# Synthetic bacterial consortium for degradation of plastic pyrolysis oil waste

**DOI:** 10.1101/2024.04.21.590079

**Authors:** Yunpu Jia, Jingxi Dou, Hendrik Ballersted, Lars M. Blank, Jianmin Xing

**Affiliations:** State Key laboratory of Petroleum Molecular & Process Engineering,, Institute of Process Engineering, Chinese Academy of Sciences, Beijing 100190, PR China; College of Chemical Engineering, University of Chinese Academy of Sciences, Beijing 100049, PR China; State Key Laboratory of Multiphase Complex Systems, Institute of Process Engineering, Chinese Academy of Sciences, Beijing 100190, China; Institute of Applied Microbiology, Aachen Biology and Biotechnology, RWTH Aachen University, Worringer Weg 1, 52074 Aachen, Germany

**Keywords:** Dearomatization, strain compounding, bacterial consortium, bioremediation, plastic degradation, neural network model, metabolic engineering

## Abstract

The plasic crisis is ominipresent, from littering macroplastic to reports that document plastic in every niche of this planet, including the human body. In order to achieve higher recycling quotas, especially of mixed plastic waste, pyrolysis seems to be a viable option. However, depending on the process parameters, plastic pyrolysis oil waste is encountered, which is difficult to valorize, due to the enormous spread of the molecules included. To reduce the molecular heterogeneity, we here artificially compounded, monitored, and optimized the performance of a bacterial consortium, which has the ability to tolerate organic pollutants and use them as energy and carbon sources for their own metabolic activity. The primary constituents of the here used plastic pyrolysis oil waste (PPOW) were alkanes and ε-caprolactam. The bacterial community exhibited noteworthy efficacy in eliminating alkanes of diverse chain lengths ranging from 71% to 100%. Additionally, within 7-days, the microbial community demonstrated a removal efficiency surpassing 50% for various aromatic hydrocarbons, along with complete eradication of ε-caprolactam and naphthalene. Besides, a back-propagation (BP) neural network method is applied to evaluate O_2_ consumption as a measure of microbial activity. The insights gained were used to build a model, which is able to predict O_2_ depletion in long-time experiments and other experimental conditions. The results are discussed in the context of a developing (open) circular plastic economy.

**Graphical abstract:** 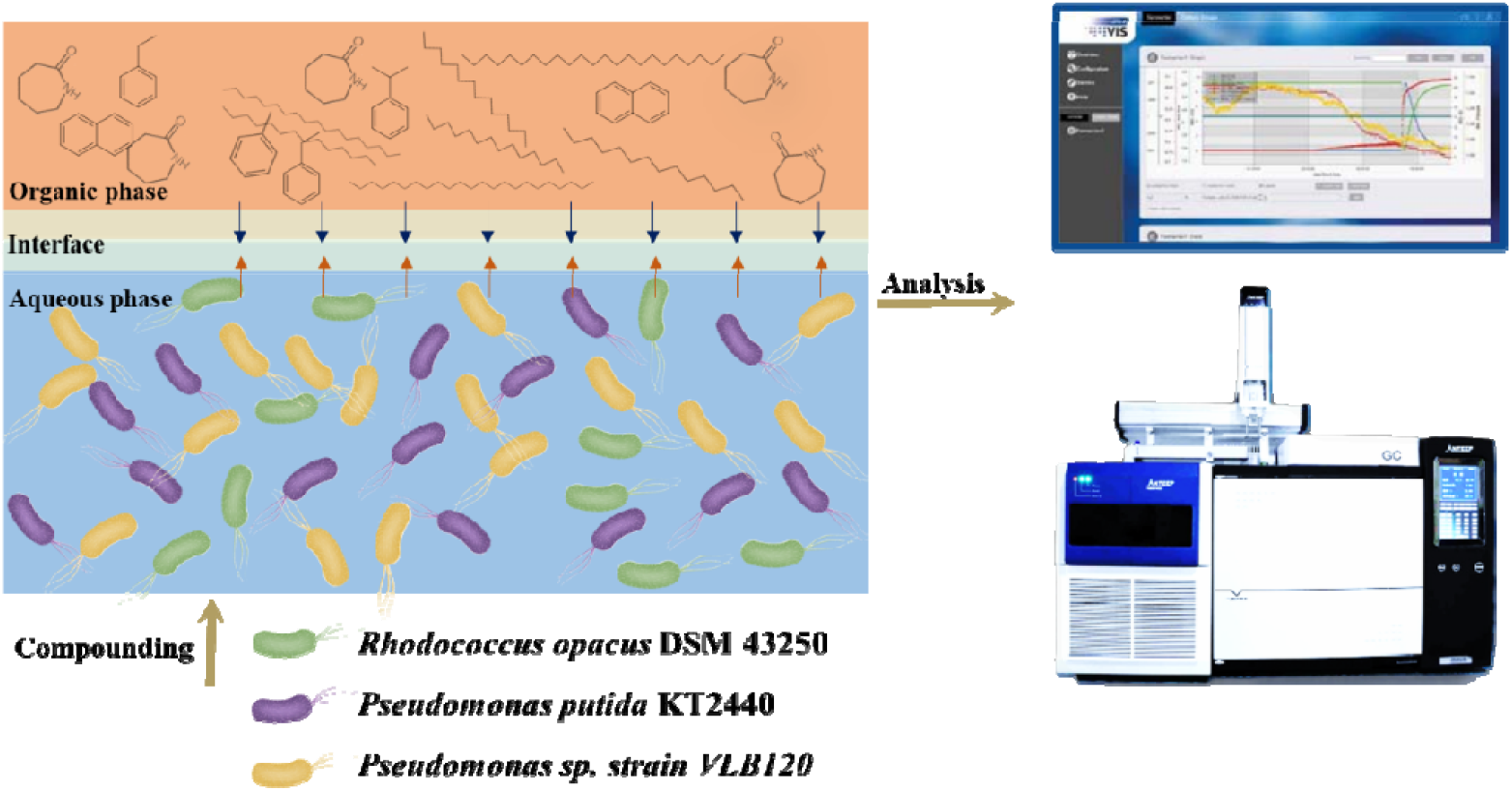

**Highlight:**

- Synthetic bacterial communities are used to remove plastic hydrolysis oil waste
- The optimized biphase reaction system can remove the majority of pollutants
- The biodegradation process can be monitored in a real-time bioprocess software
- Neural network techniques are used to model and predict the removal process

## 1. Introduction

Plastics have become the most relevant materials in today’s societies. Worldwide plastic production is anticipated to double within the next two decades and is projected to nearly quadruple by 2050 [1]. Besides mechanical recycling being limited, non-recovered plastics cause severe pollution and threaten life of animals and humans. The necessity for a “post-consumer plastics economy” has become evident and is influencing political and business choices [2]. The key to achieving a circular economy is to identify appropriate technologies. Reprocessing plastics without compromising function and value necessitates a paradigm shift in our understanding of plastic recycling. The hydrocarbon resource targeted for recovery has to demonstrate potential for diverse and high-value applications [3], also coined open loop-recycling. This ranges from plastics’ conversion into highly efficient fuels to input oil products for the polymer industries.

CARBOLIQ GmbH (Remscheid, Germany) has proven to transform mixed plastic waste into a liquid resource by applying a one-stage conversion technology, catalytic tribochemical conversion (CTC), a direct liquefaction process combined application of thermal, catalytic, and mechanochemical (tribochemical) mechanisms. CTC works in moderate conditions where atmospheric pressure and process temperatures are around 400 °C. The solid input material is continuously fed to the hot liquid system enriched with a catalyst. The oiled plastic is separated by destillation, with the result of a plastic pyrolysis oil waste (PPOW) fraction. This fraction might be difficult to valorize, as it constitued a mixture of molecules with different chemical and physical protperties.

Considering the complexity of the substrate, we propose a strategy to treat this complex fraction using artificially compounded strains in a synthetic microbial consortium. *Pseudomonads* play a crucial role in element cycling [6] and exhibit significant potential for bioremediation. They boast a diverse repertoire of pathways for degrading various non-natural and non-inherent pollutants, particularly aromatic organics [7–10]. Among these, *Pseudomonas putida* KT2440 [11,12] stands out as the most extensively characterized saprophytic *Pseudomonas*, maintaining its ability to thrive and perform essential functions in the environment. It has the catabolic potential against a wide range of natural aromatic compounds. *Pseudomonas spec*. VLB120 [13,26] underwent examination for its inherent tolerance to toxic compounds like toluene and styrene. At the genomic level, homologs of known solvent-tolerance genes were investigated. *Rhodococcus* bacteria are widely acknowledged for their significant potential in bioremediation and have been successfully employed for the removal of contaminants from soil, water, and air [14]. *Rhodococcus opacus* DSM 43250 is a gram-positive bacteria that was first isolated from soil. Prominent metabolic characteristics encompass limited catabolic repression by more readily available carbon sources [15] and adaptable biodegradation pathways for both sugars and aromatics [16,17].

In this study, the microbial consortium consisted of three bacterial strains, *Rhodococcus opacus* DSM 43250, *Pseudomonas putida* KT2440, and *Pseudomonas spec*. VLB120. The PPOW was characterized and used as carbon and energy source for the synthetic microbial consortium. The degradation of PPOW by bacterial consortium was evaluated after process enhancement for the degradation ratios of each of the major components it contains. For growth charaterization of the consortium, a three-layer back-propagation (BP) neural network was used to represent the experimental data and predict oxygen consumption in further experiments. The results of the study are discussed in the context of PPOW as a carbon and energy source for bacterial consortium for the purpose of contained organic components removal.

## 2. Materials and methods

### 2.1 Chemicals

PPOW was supplied by CARBOLIQ GmbH (Remscheid, Germany). Alkane standard solution was purchased from Merck (Darmstadt, Germany). Water was purified using a Milli-Q Academic System (18.2 MΩ cm; 0.22-μm filter) (Millipore, Molsheim, France). Other chemicals used in this work were obtained from Carl Roth (Karlsruhe, Germany) and Sigma-Aldrich (St. Louis, MO, USA), unless stated otherwise.

### 2.2 Components of culture medium

The precultures were made in the Delft mineral salt medium (MSM). The final medium composition per liter was 3.88 g of K_2_HPO_4_, 1.63 g of NaH_2_PO_4_, 2.00 g of (NH_4_)_2_SO_4_, 0.1 g of MgCl_2_· 6 H_2_O, 10 mg of EDTA, 2 mg of ZnSO_4_ · 7 H_2_O, 1 mg of CaCl_2_ · 2 H_2_O, 5 mg of FeSO_4_ · 7 H_2_O, 0.2 mg of Na_2_MoO_4_· 2 H_2_O, 0.2 mg of CuSO_4_ · 5 H_2_O, 0.4 mg of CoCl_2_ · 6H_2_O, and 1 mg of MnCl_2_ · 2 H_2_O, supplemented with 20 mM sodium benzoate as carbon source. Precultures of single strains consisted of 10 mL medium in 50 mL Erlenmeyer flasks. The individual cultures were shaken at 200 rpm and were incubated at 30 °C.

### 2.3 Bacterium

*Rhodococcus opacus* DSM 43250, *Pseudomonas putida* KT2440, and *Pseudomonas spec*. VLB120 were stored in the strain collection of the Institute of Applied Microbiology (RWTH Aachen University, Aachen, Germany).

### 2.4 Biodegradation reaction

For the two-phase reaction, cells were cultured at the specified temperature in a modified MSM [18]. The medium’s standard phosphate buffer capacity was increased three-fold to reduce the influence of acid during the growing progress of *Pseudomonas*. The final aqueous culture medium composition per liter was: 11.64 g of K_2_HPO_4_, 4.89 g of NaH_2_PO_4_, 2.00 g of (NH_4_)_2_SO_4_, 0.1 g of MgCl_2_ · 6H_2_O, 10 mg of EDTA, 2 mg of ZnSO_4_ · 7H_2_O, 1 mg of CaCl_2_ · 2H_2_O, 5 mg of FeSO_4_ · 7H_2_O, 0.2 mg of Na_2_MoO_4_ · 2H_2_O, 0.2 mg of CuSO_4_ · 5H_2_O, 0.4 mg of CoCl_2_ · 6H_2_O, and 1 mg of MnCl_2_ · 2H_2_O. Ethyl decanoate or 2-undecanone was selected as the organic phase in the reaction system, which contains plastic pyrolysis oil waste as the main carbon and energy source, and 0.05% (wt%) yeast exact was supplied to promote the growth of the strains. To explore the O_2_ consumption rate under various cultivation conditions, 200 mL anaerobic media bottles with aluminum seals were employed. The total volume of the biphasic medium was 20 mL. Aeration was only from the head space in the bottles. The cell densities of the individual strains were quantified using the spectrophotometer determined at OD_600_. The aforementioned experimental conditions were also employed for experiments measuring CO_2_ production, with the only difference being that the reaction volume was 50 mL medium in a 1 L Schott-Duran bottle.

### 2.5 Extraction of residual PPOW

Residual PPOW was sampled by liquid-liquid extraction. Briefly, broth culture (50 mL) was extracted twice with a five-volumes of n-hexane. After removing the aqueous phase with a separating funnel, the residual concentration of PPOW was determined by gas chromatography combined with a flame ionization detector (GC-FID).

### 2.6 Analytical methods

#### 2.6.1 CO_2_ measurement

BCP-CO_2_ gas analyzer in combination with a bioprocess software (BlueSens Gas Sensor GmbH, Herten, Germany) was used for in-situ, continuous, and online real-time CO_2_ analysis. The CO_2_ concentration was tracked using infrared light, attenuated by the analyte gas, and reflected into the sensor’s detector unit. The sensor was airtightly affixed to the opening of a 1 L static Schott-Duran bottle used as the culture vessel. Measurements were conducted without air exchange for a period of up to 1 week, with the longer runtimes constrained by the availability of oxygen for microbial metabolism.

#### 2.6.2 Gas-chromatography to assess O_2_ depletion

The O_2_ was measured by an SRI 8610C Multi-detector Gas Chromatograph (GC) coupled with a helium ionization detector (HID). HayeSepD Pre-column (2 mm × 2 m) and main column (Molsieve 13x) were used to separate O_2_ from samples containing a mixture of gases. O_2_ content is obtained according to the standard curve. The column was connected to a thermal conductivity detector (TCD, 157 °C) at higher concentrations and a HID (100 V, 204 °C) at lower concentrations. A 48 mL/min helium gas flow was applied, and the column was operated isothermally at 60 °C.

#### 2.6.3 Gas chromatography-mass spectrometry (GC-MS)

The original sample and alkane standard solution were analyzed using GC-MS after filtration and a tenfold dilution with n-hexane. The GC-MS analysis was performed using a GC-MS QP2010 Ulltra, SHIMADZU and 30 m long Rtx-5MS column (internal diameter, 0.25 mm, film thickness, 0.25 μm). Helium served as the carrier gas with a flow rate of 3 mL/min, and 1 μL of the sample was injected into the GC. The GC oven temperature was programmed to start at 50 °C, increasing by 6 °C/min up to 280 °C, and held for 5 minutes. The ion source was set at 200 °C.

#### 2.6.4 GC-FID

Hexane extracts (1.0 μL) of PPOW were analyzed with Hewlett Packard 5890 Series II GC equipped with a flame ionization detector (FID) and 30 m long ZB-WAX column (internal diameter, 0.25 mm, film thickness, 0.25 μm). The carrier gas was nitrogen. The injector and detector temperatures were maintained at 200 and 290 °C, respectively. The GC column was programmed at an initial temperature of 40 °C and held for 1 minute. It was then increased by 1 °C/min up to 50 °C, followed by a rapid ramp at 40 °C/min to 200 °C, and finally stopped running at 35 minutes.

## 3. Results and discussion

### 3.1. Component of PPOW

GC-MS and GC-FID were used to determine the composition of the PPOW. Given that the end product of the plastic pyrolysis process is alkanes of various chain lengths, it is hypothesized that the sample should encompass a substantial quantity of alkanes. It is also known from the raw material of the product that the sample should also contain various aromatic compounds. From the result of GC-MS in **Fig. 1**, the predominant peaks in the sample were identified as alkanes, and the mass spectrometry results revealed that the elevated content of substances other than alkanes comprised cumene, toluol, ethylbenzol, naphthalin, and ε-caprolactam. The highest content was of ε-caprolactam (8032 μg/mL) even far exceeding that of alkanes. We verified this result with GC-FID and determined the content of these five substances (**Fig. S1, S2 and Table. S1)**.

**Fig. 1.**
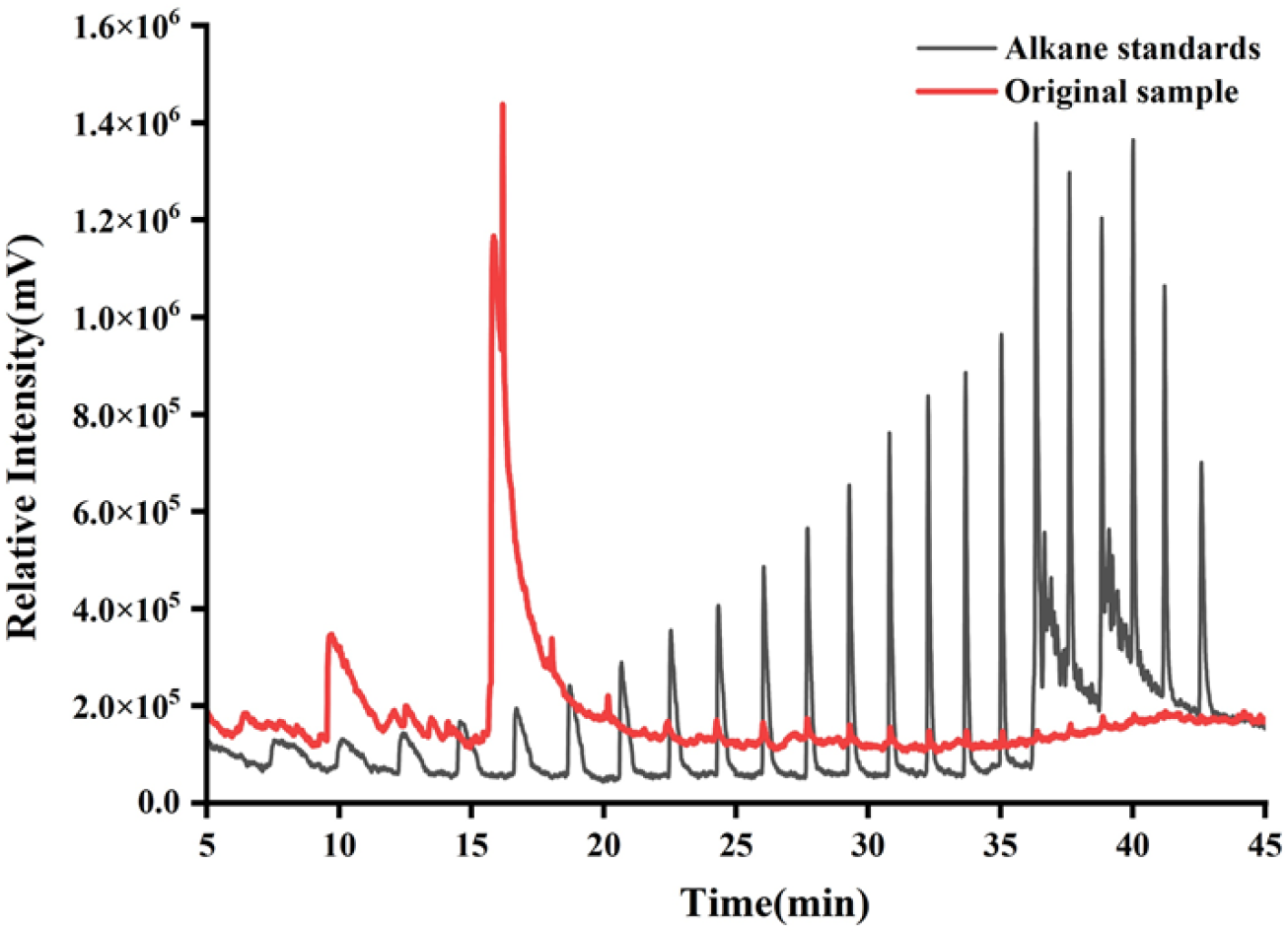
Substrate characterization by GC-MS. The plastic pyrolysis oil waste (PPOW, red) was filtered and diluted 5 times by n-hexane and was compared against an alkane standard.

### 3.2. Intensification of biodegradation of PPOW in organic-aqueous two-phase system

Biodegradation of PPOW by synthetic bacterial communities comprised of *Rhodococcus opacus* DSM 43250, *Pseudomonas putida* KT2440 and *Pseudomonas spec*. VLB120 were investigated in an organic-aqueous two-phase system. To mitigate the toxic effects of PPOW on strains, previouly used second phase compounds, namely 2-nonanone [24]and ethyl decanoate [18], served as a second organic phase each. Their role was to reduce the concentration of the organic components in the water phase, while having a substrate reservoir. The BCP-CO_2_ gas analyzers were used for online real-time CO_2_ analysis [19]. The final second phase was determined by comparing the release of CO_2_ from the two-phase reaction system. When using 2-nonanone as the second phase (**Fig. 2**), CO_2_ exhibited a significantly higher release in a shorter period compared to the culture with ethyl decanoate. This suggests that when utilizing 2-nonanone as the second phase, the bacterial communities exhibits a higher metabolic activity, and was chosen here for the further experiments.

**Fig. 2.**
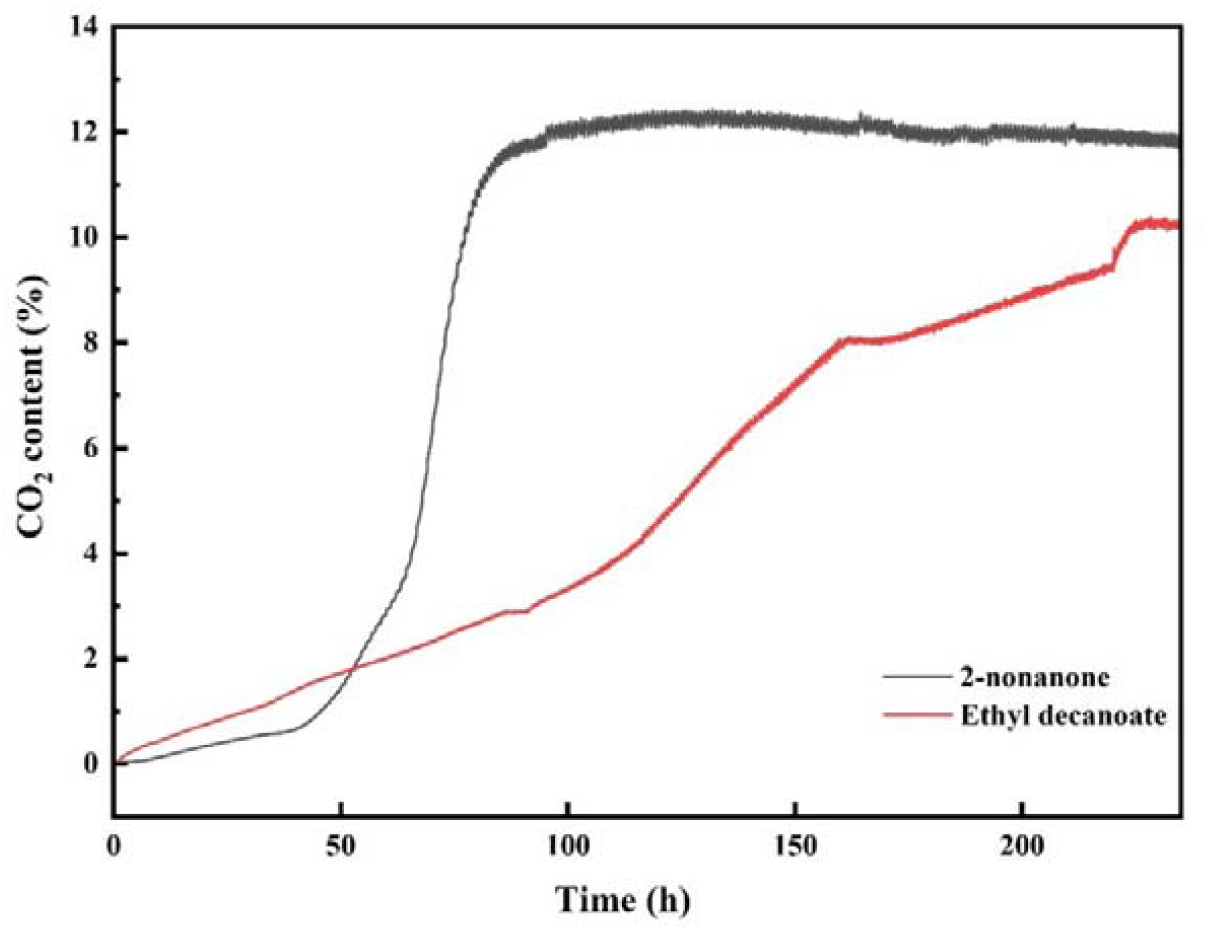
Comparison of the second organic phase compounds. The suitability of 2-nonanone and ethal decanoate as secon phase was evaluated using CO_2_ measurements. The CO_2_ released was online measured during the biodegradation of PPOW in an oil-water two-phase system. The conditions were 30 □, with a second phase-water ratio of 1:5, a PPOW content (%V) of 20% in the second phase, and a strain compounding ratio of *Rhodococcus opacus* DSM 43250: *Pseudomonas putida* KT2440: *Pseudomonas spec*. VLB120 was 1:1:1.

Afterward, we investigated other factors influencing the metabolic activity of the bacterial consortium and determined the optimal culture conditions based on these findings (**Fig. 3**). The optimal growth temperature for the three strains in the bacterial consortium was found to be 30 L. However, it is crucial to note that the optimum temperature for the growth and metabolism of strains is typically not uniform [11,20]. Some microorganisms may display a faster growth rate at lower temperatures, yet their metabolic activity could be more pronounced at higher temperatures. In addition to monitoring the CO_2_ release rate, assessing O_2_ consumption is equally vital as a key indicator of microbial metabolic activity [21–23]. Therefore, to simplify the experiments, all optimization trials measured the O_2_ consumption at different time points under dynamic shaking flask cultivation conditions. Results in **Fig. 3a** indicate that the optimal temperature for the bacterial consortium’s utilization of organic substances in PPOW aligns with the growth temperature. Extreme temperatures, whether high or low, prove unfavorable for the absorption and utilization of carbon sources in PPOW. This could be attributed to the temperature’s impact on the metabolic activity of enzymes within the bacterial consortium. Additionally, temperature can influence the solubility of O_2_ in the biphasic reaction solution, thereby directly affecting the microbial utilization of O_2_. In a biphasic reaction system, the ratio of the organic phase to the aqueous phase is undoubtedly a crucial factor influencing the reaction rate by affecting concentrations of organic matter exposed to the bacterial consortium. Maximum degradation was achieved in the 1:5 organic: aqueous ratio. Further increasing the ratio did not lead to an increased degradation rate. Therefore, a 1:5 ratio was selected for subsequent studies. Excessive concentrations of PPOW can exert a toxic effect on the microbial community, while lower concentrations may not fully exploit the degradative capabilities of the bacterial consortium. The findings suggest that when PPOW constitutes 20% of the second phase, O_2_ depletion in the closed system occurs most rapidly, implying optimal removal efficiency (**Fig. 3c**).

**Fig. 3.**
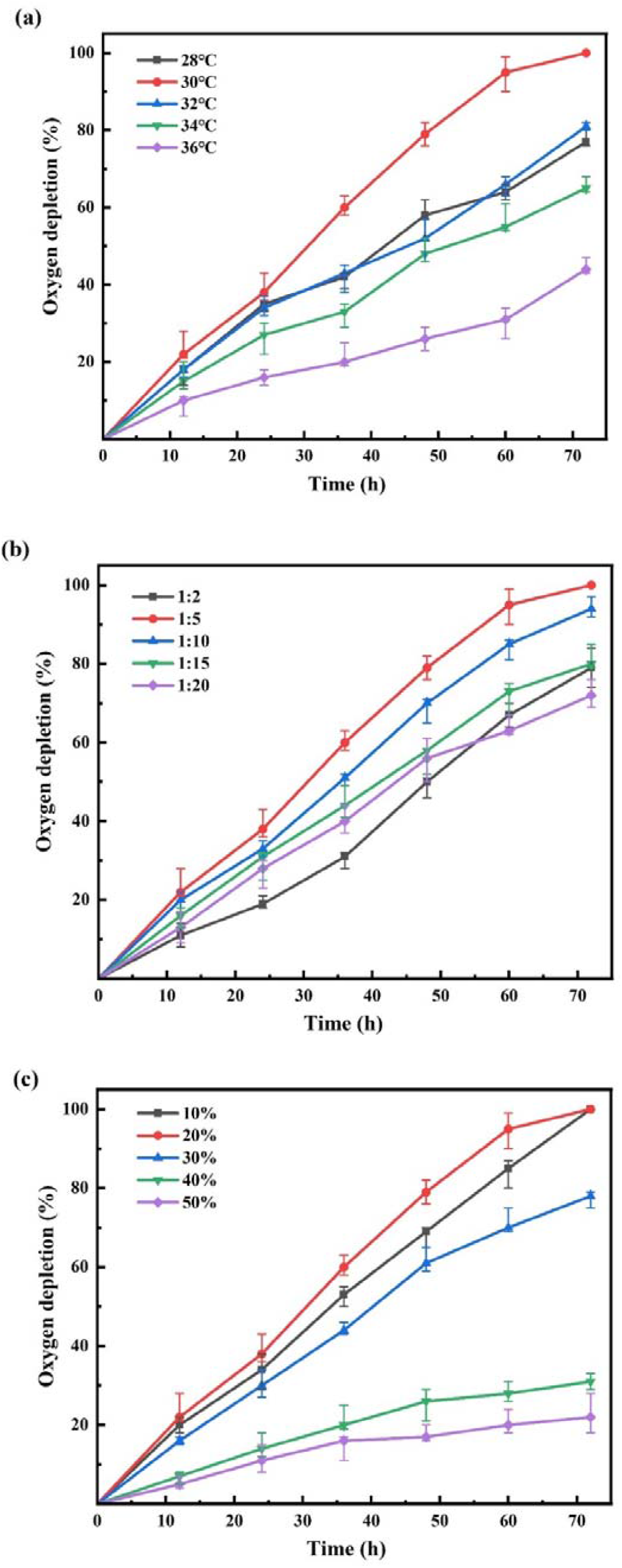
The rate of O_2_ depletion under different growth conditions of the synthetic consortium. (a) temperature, (b) organic:aqueous phase ratio, and (c) PPOW content (%V) in the second phase. The strains compounding ratios of *Rhodococcus opacus* DSM 43250: *Pseudomonas putida* KT2440: *Pseudomonas spec*. VLB120 was 1:1:1.

In addition, we explored the impact of varying strain compositions within bacterial consortium on the removal of PPOW (**Fig. 4**). Differing from the previous experimental conditions, the microbial community displayed maximum metabolic activity when the ratio of *R. opacus* DSM 43250 to *P. putida* KT2440 to *Pseudomonas* spec. VLB120 was 1:1:2. This specific composition of the bacterial consortium exhibited the quickest CO_2_ production in the reaction system. However, the specific mechanism remains unclear, and it is speculated that the enhanced tolerance of *Pseudomonas* spec. VLB120 to toxic components may play a role [25].

**Fig. 4.**
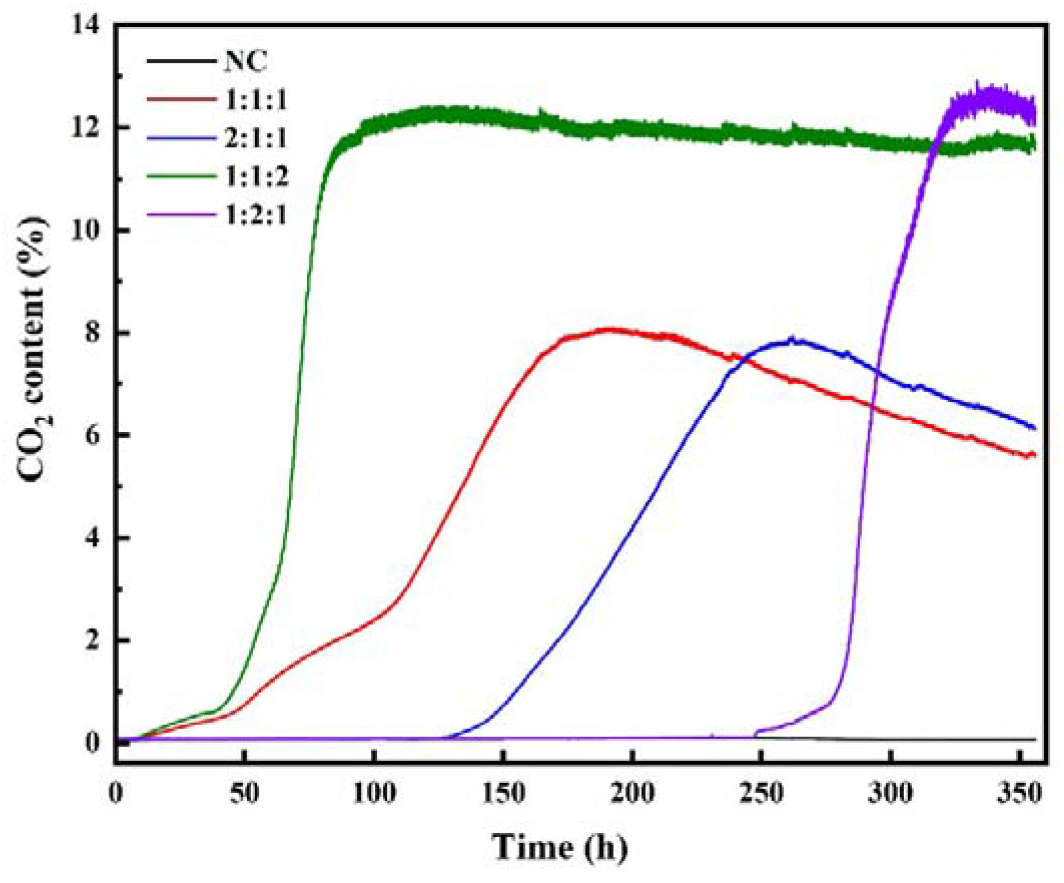
Evaluating bacterial consortium composition. The rate of CO_2_ release varies with different strains compounding ratios of *Rhodococcus opacus* DSM 43250: *Pseudomonas putida* KT2440: *Pseudomonas spec*. VLB120. NC: negative control groups without adding any bacteria.

### 3.3. Removal ratio of components in PPOW

Therefore, the subsequent long-term studies were carried out for the degradation of aromatic and aliphatic compounds present in PPOW at 30 □, 1:5 ratio of organic: aqueous phase, 1:1:2 ratio of *R. opacus* DSM 43250: *P. putida* KT2440: *Pseudomonas spec*. VLB120 and 20% PPOW content (%V) in the organic phase. GC-FID was employed to detect changes in PPOW before and after degradation, and to calculate the removal rate (**Fig. 5**). Percentage degradation of several key components present in the PPOW after 7 days by bacterial consortium in the biphasic reaction system is shown in **Fig. 6**. With the exception of toluol, the removal rates for more than a dozen substances with relatively high concentrations in the waste all surpass 50%, with some achieving complete removal after 7 days of biotreatment by the snthetic bacterial communities. However, it is currently unclear what the metabolic products are. After microbial treatment, a distinct alteration in the color of the organic phase in the two-phase system is evident when compared to its pre-treatment state **(Fig. S4)**. This color transformation serves as a clear indication of the removal of organic substances in the treated PPOW.

**Fig. 5.**
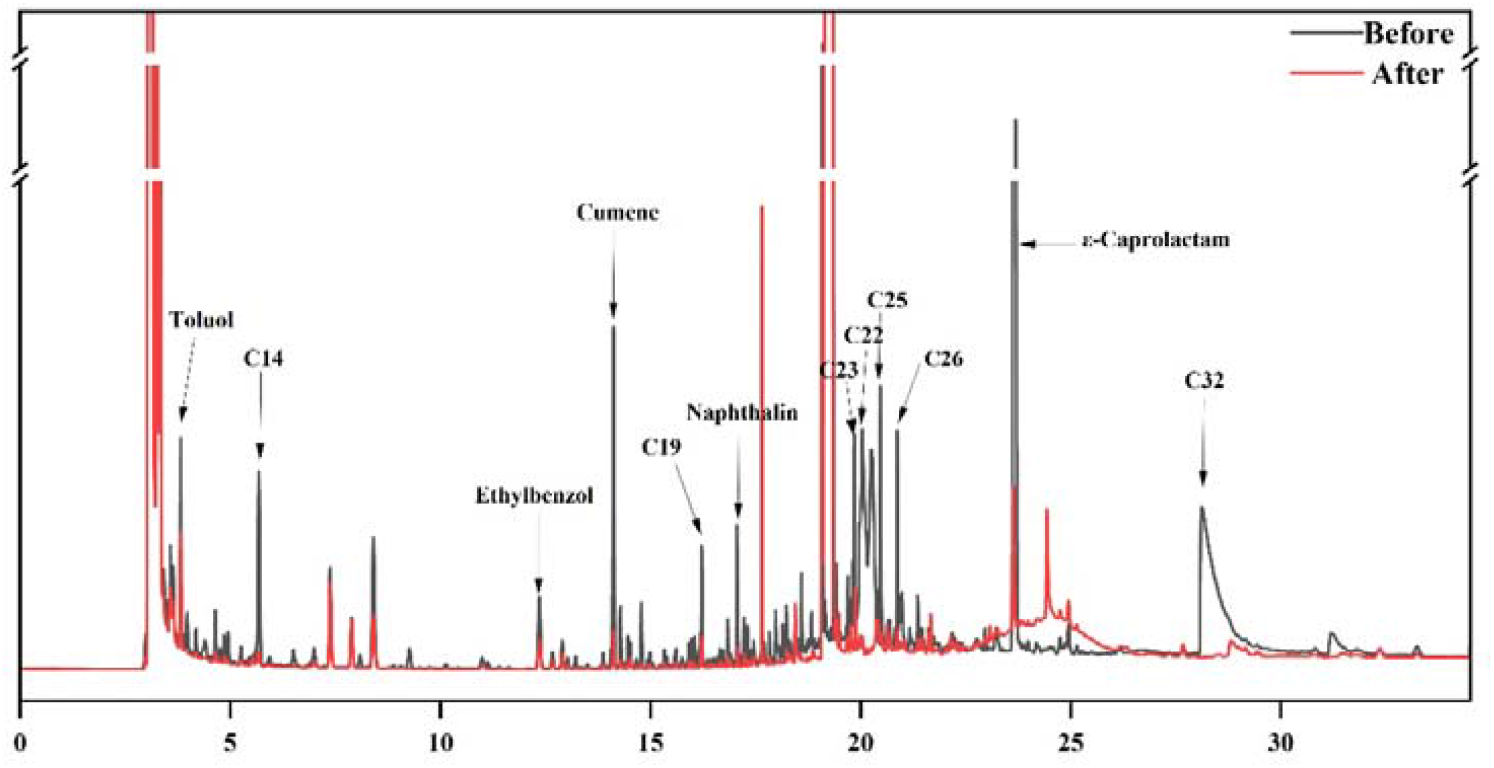
Investigating substrate uptake. GC-FID chromatograms of original PPOW (black) and PPOW used as carbon and energy source by the synthetic bacterial consortium under optimal conditions (red) after 7 days.

**Fig. 6.**
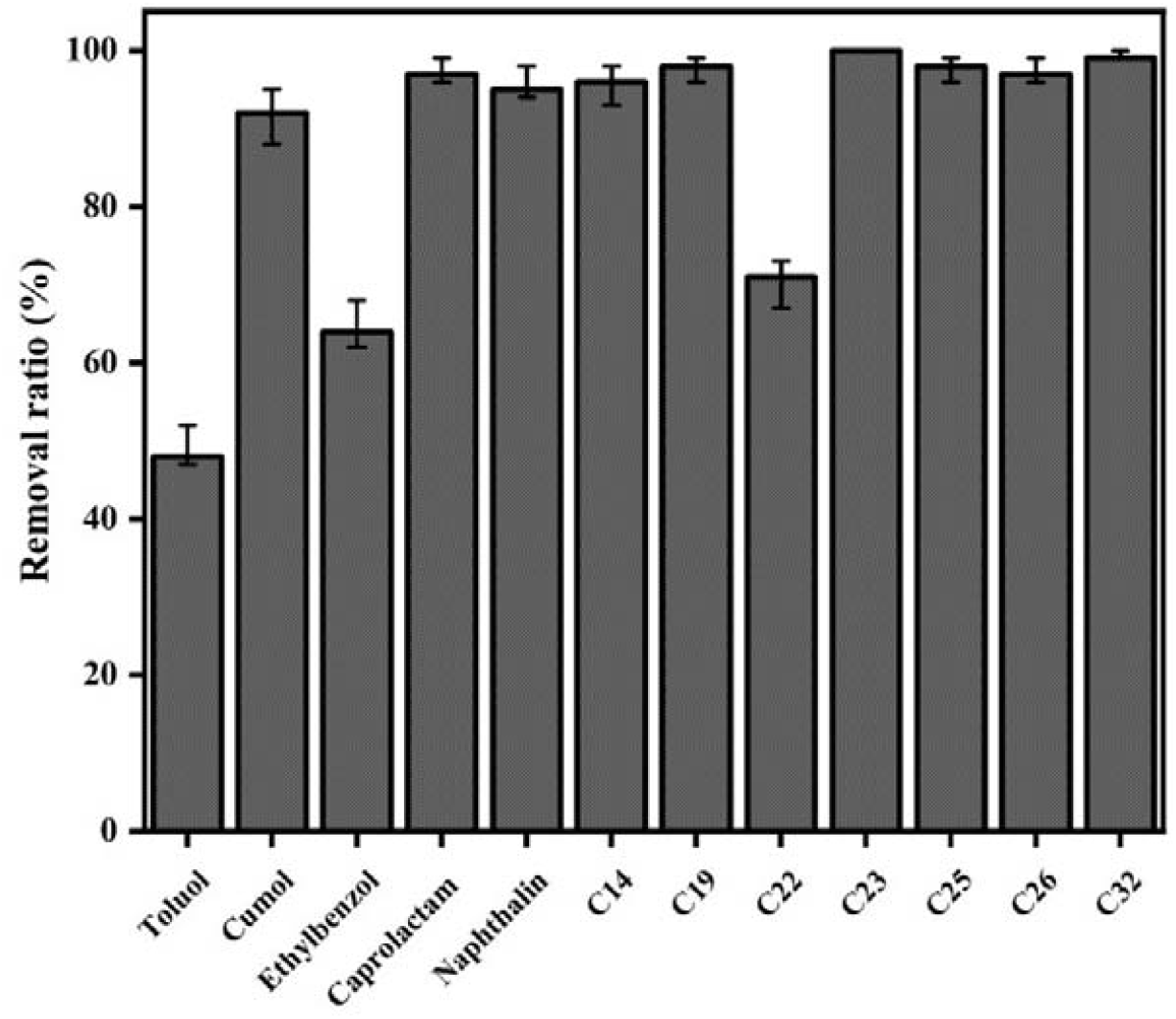
Investigating substrate uptake. The removal ratio of major substances from PPOW under optimal conditions after 7 days. Each error bar represents the standard deviation of three replicate experiments. Removal ratio□= □(Initial concentration□-□Final concentration) □×□ 100/ Initial concentration.

### 3.4. Neural Network Model for O_2_ depletion in organic-aqueous two-phase system

As oxygen availibility is key for PPOW activation and degradation, we investigated the data using an AI model. A three-layer Back-propagation (BP) neural network, which is a multilayer feed-forward neural network, with enough nodes in the hidden layer has the ability to simulate any complex nonlinear mappings, hence has strong fitting ability. Therefore, a BP neural network was used to represent the experimental data and to predict oxygen consumption in further experiments. The model is written as

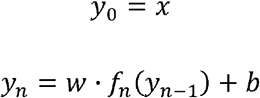

where *x* is the input variable, defined as time and influence factors (temperatures, organic: aqueous phase ratio and PPOW content (%V) in the second phase), and *y*_*n*_ is our excepted prediction (O_2_ depletion). *f*_*n*_ is the activation function in *n* layer. *w* and *b*are the parameters of the neural network, namely weight and bias, which result from the training. Many optimization algorithms could be used in the training process, while here the Levenberg-Marquardt (LM) algorithm was chosen (**Table 1**).

**Table 1.**
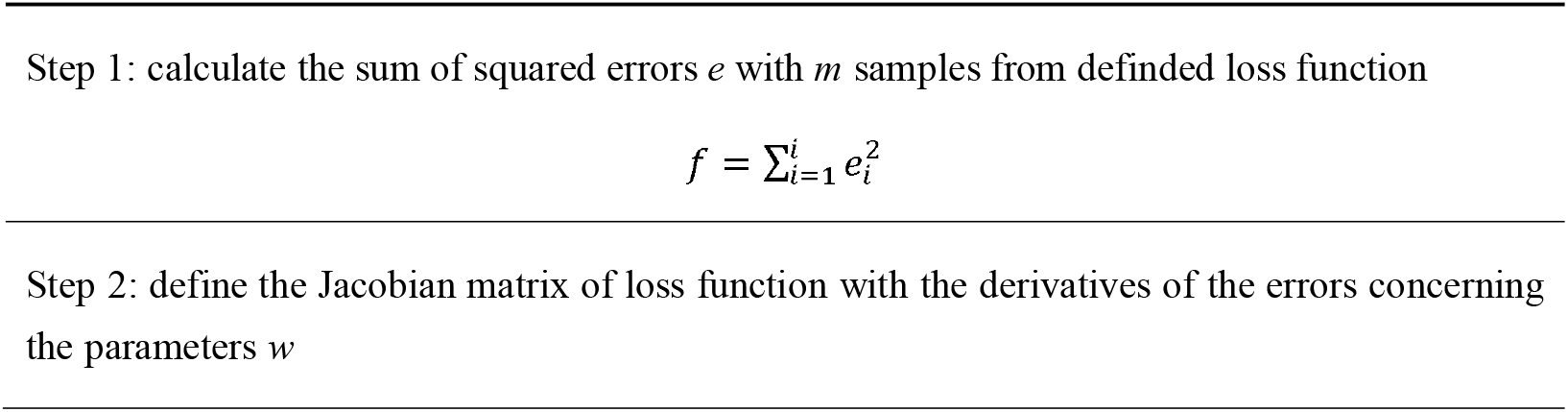
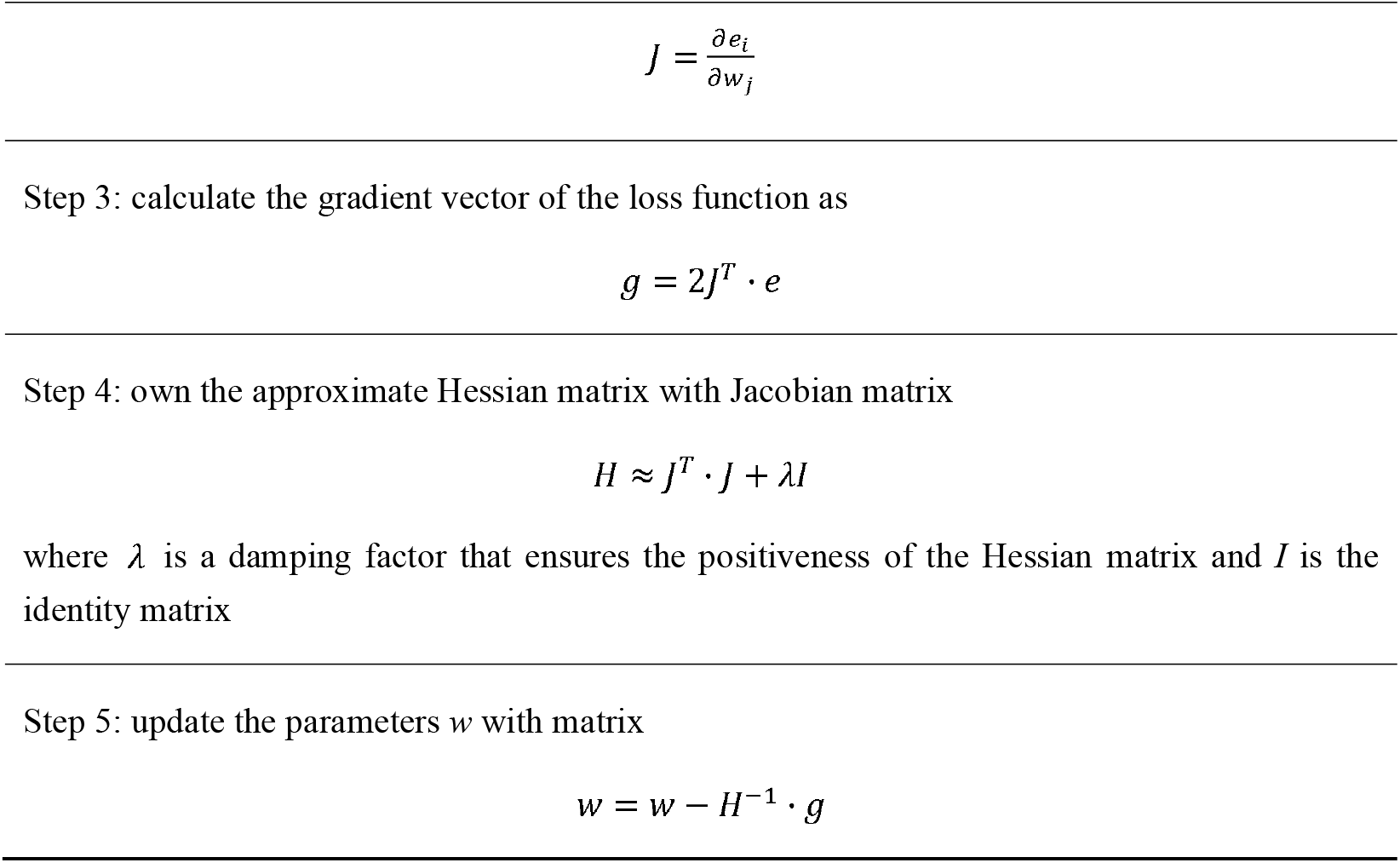
Levenberg-Marquardt (LM) optimization algorithm.

Three BP neural networks were built and trained. Their layers, weights and bias are listed in **Table 2**. Firstly, the input influence factors was time and temperatures, and the output prediction was O_2_ depletion. The mean square error (RMSE) and the correlation coefficient *R*^*2*^ are values to evaluate model performance, written as

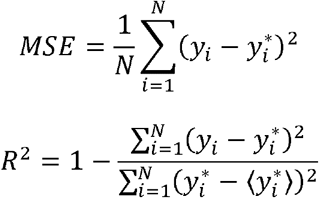

where 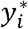 is the experimental data, or expected data, *y*_*i*_ is the predicted value from the neural network model, and 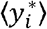 is the average experimental data. The BP neural network exhibits well fitting results, with an MSE of 2.06×10^−3^ and an *R*^*2*^ of 0.9903. As discussed above, the best results are obtained at 30 °C (**Fig. 3a**). Results from the BP neural network satisfy and suggest a possible range of 29 to 32 °C (**Fig. 7a**). With this model, the degradation time of PPOW by the synthetic bacterial consortium can be predicted well. For example, after 4 days more than 80% O_2_ is depleted, and almost all O_2_ could be depleted after 6 days.

**Table 2.**
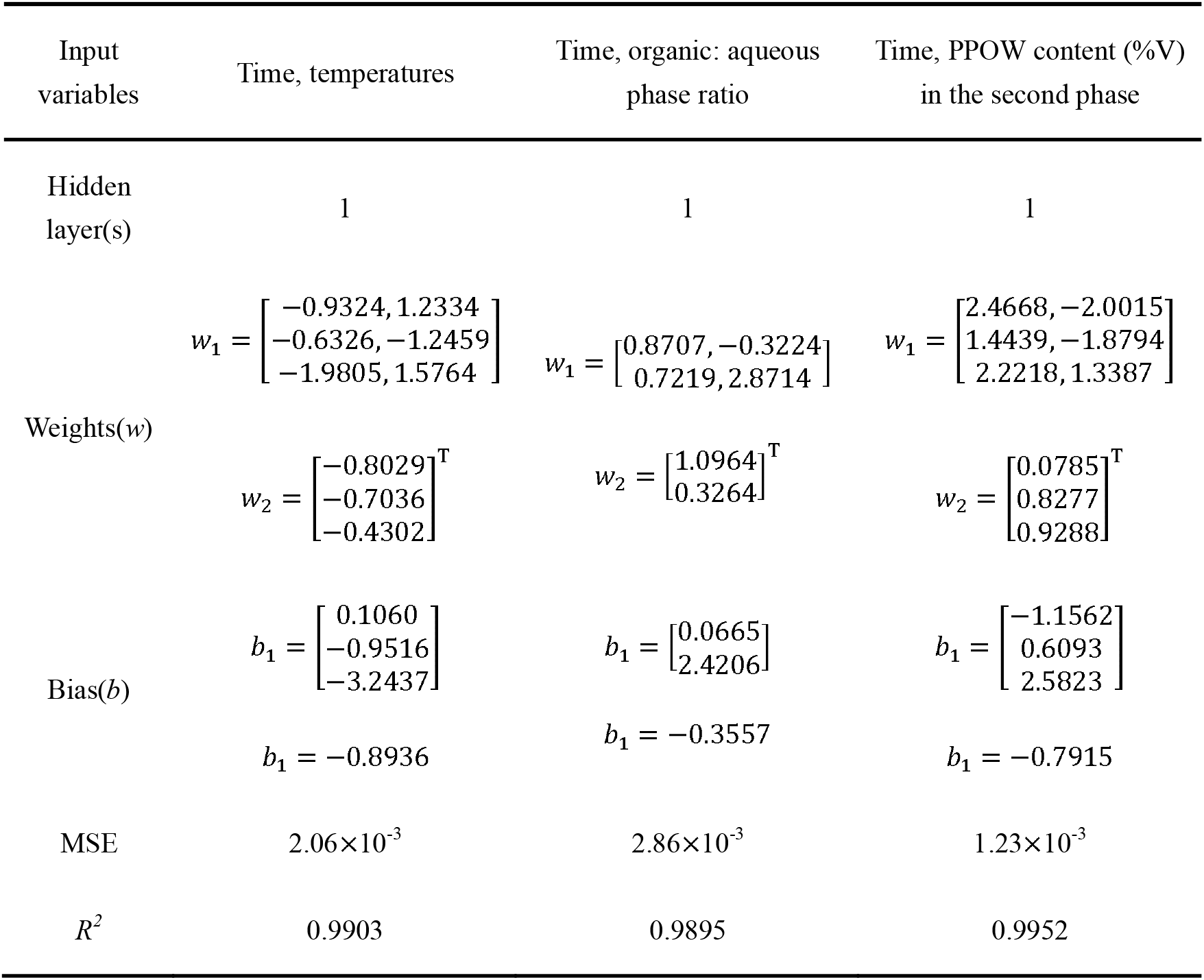
BP neural parameters with output variable O_2_ depletion.

**Fig. 7.**
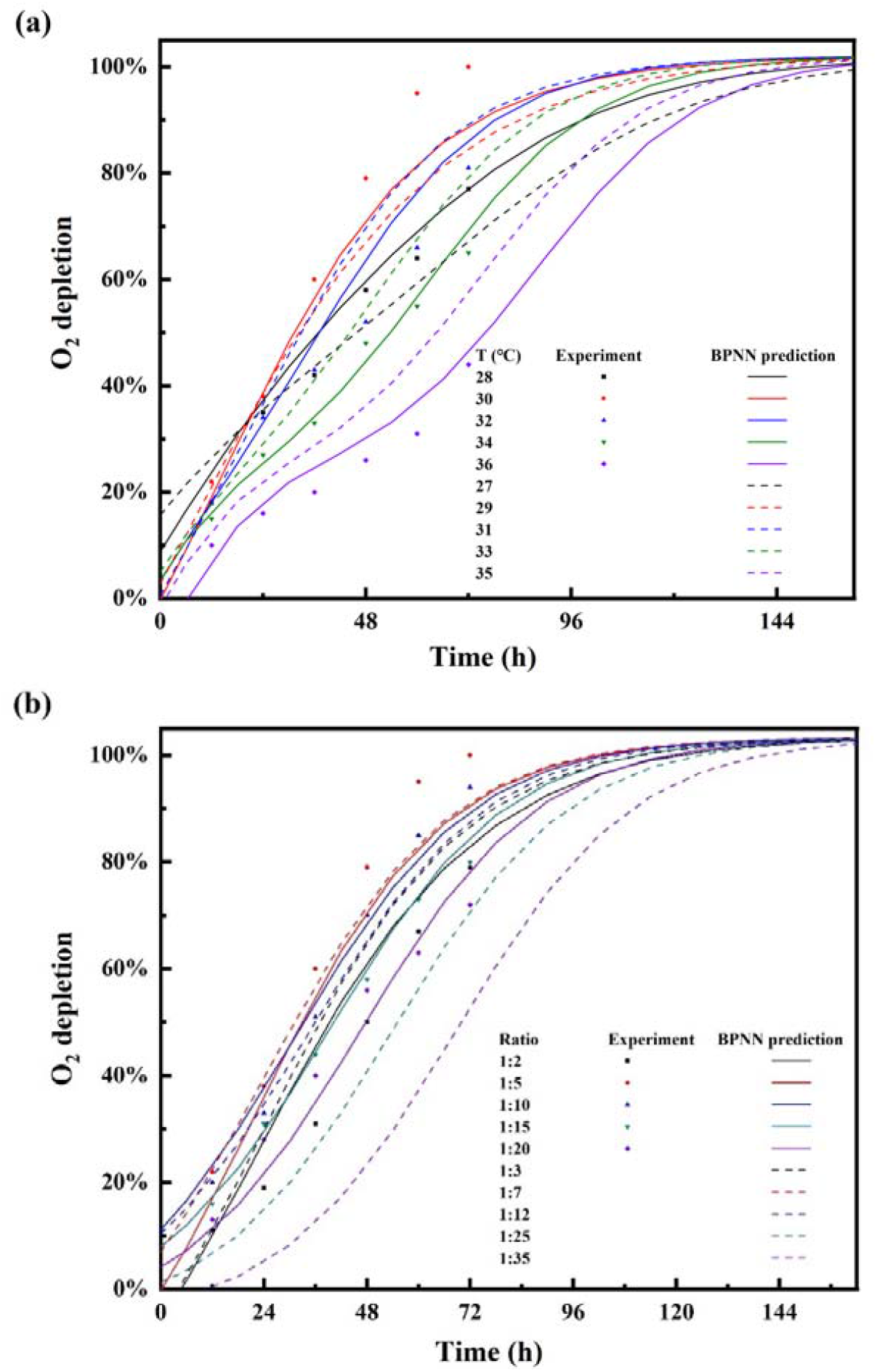
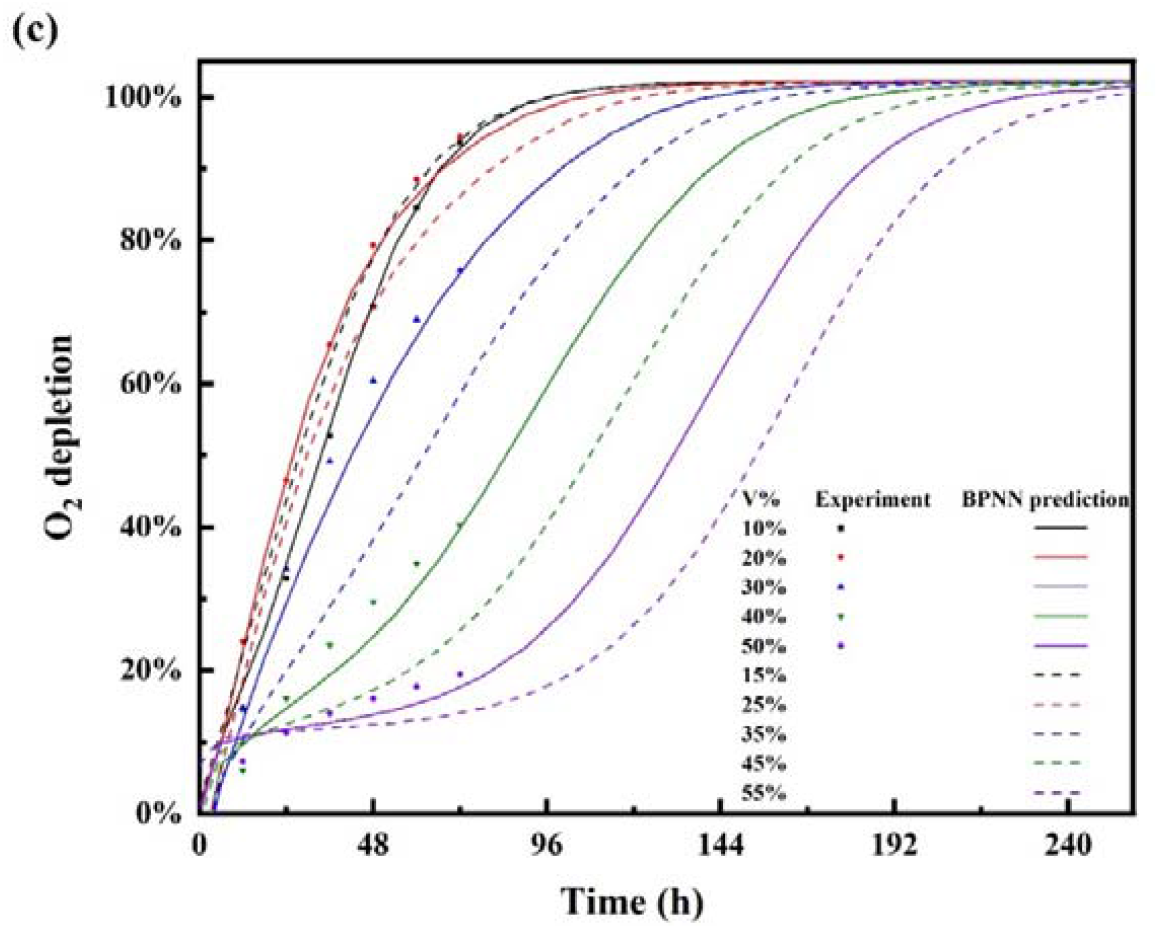
Fitting and predicting oxygen depletion in synthetic consortium cultures with PPOW as substrate. Oxygen depletion was estimated using the BP neural network models under different parameter influences: (a) temperature, (b) organic:aqueous phase ratio, and (c) PPOW content (%V) in the second phase (c).

Secondly, when the input influence factors are time and organic:aqueous phase ratio, oxygen depletion was also well predicted by the BP neural network model (**Fig. 7**). The MSE was 2.86×10^−3^ and the *R*^*2*^ was 0.9895. Although some discrete data points are far from the neural network curve (ratio-1:5), the predicted results still fit the experimal data (**Fig. 3**). Besides, the neural network model even forecast O_2_ depletion in time (T=0∼162 h) and other phase ratios (1:3, 1:7, 1:12, 1:25, 1:35). The model suggests that the higher the phase ratio, the slower the O_2_ depletes. For example, it is predicted that after 4 days more than 80% O_2_ could be depleted with an organic:aqueous phase ratio lower than 35%. However, it should be noticed that the BP neural network model has a poor predictability at the start of the cultivation (before 0.5 days). It can be explained by our principle of parameter optimization, that the model results must converge to 1 at later times. Therefore, in the training process the limited network structure and experimental replicates do not allow a better prediction to describe the early time. More high quality experimental data would be required to improve the model.

Thirdly, with the input influence factors time and PPOW content (%V) in the second phase, the BP neural network model shows results in time (T=0∼258h) and PPOW content (%V=15%, 25%, 35%, 45%, 55%) (**Fig. 7c**). The BP neural network exhibits good prediction, with an MSE of 1.23×10^−3^ and an *R*^*2*^ of 0.9952. The results confirm a PPOW content of 20% of the second phase, and expose when the PPOW content (%V) is 15%∼25%, the degradative capabilities of the synthetic bacterial consortium can be exploited well. In addition, oxygen depletion can be predicted with the neural network, suggesting after 5 days more than 80% O_2_ is depleted with a PPOW lower than 35%, and after 10 days amlost all O_2_ is consumed with a PPOW lower than 55%. The predictability appears to increase with incubation time, correlating with the number of data available.

In summary, three BP neural network modesl were built based on experiment data. We aimed to describe the tendency of O_2_ depletion with different factors influencing the outcome (temperatures, organic:aqueous phase ratio, and PPOW content (%V) in the second phase). These BP neural network models are able to forecast the time required for PPOW degradation using the presented artificial bacterial consortium. As always with AI, more experimal data would be helpful to improve the models in the future.

## Conclusions

By skillfully combining a bacterial consortium with high degradation capability for plastic pyrolysis oil waste within a two-phase system, we contribute a promising solution for end-of-life plastic treatment that is complementary to the many other suggestions. This approach not only addresses plastic resource utilization but also contributes to the advancement of a bio-circular economy within the domain of plastics. The synthetic bacterial communities showed excellent ability in degrading PPOW. Besides, a BP neural network method is applied to evaluate O_2_ consumption. The model presented is able to predict O_2_ depletion at long cultivation times and can extrapolate to other experimental conditions. The bacterial community can efficiently remove alkanes of varying chain lengths. Further studies aimed at understanding the mechanisms of synergistic interactions among the bacterial consortium and the identification, as well as the upgrading and reconstruction of metabolic products, need to be conducted in the near future.

## Acknowledgments

The authors gratefully acknowledge the financial support provided by the National Natural Science Foundation of China (grant numbers 31961133017, 31961133018, 31961133019). These grants are part of “MIXed plastics biodegradation and UPcycling using microbial communities” MIX-UP research project, which is a joint NSFC and EU H2020 collaboration. In Europe, MIX-UP has received funding from the European Union’s Horizon 2020 research and innovation program under grant agreement No 870294.

LMB acknowledges support of the Werner Siemens Foundation in the frame of the WSS Research Centre “Catalaix”.

## CRediT authorship contribution statement

Yunpu Jia: Conceptualization, Methodology, Validation, Data curation, Writing-original draft, Writing-review & editing. Jingxi Dou: Methodology, Validation, Data curation. Hendrik Ballersted: Methodology. Lars M. Blank: Conceptualization, Writing-review & editing, Supervision, Project administration. Jianmin Xing: Conceptualization, Writing-review & editing, Supervision, Project administration.

## Declaration of Competing Interest

The authors declare that they have no known competing financial interests or personal relationships that could have appeared to influence the work reported in this paper.[1][1] M.W. Guzik, T. Nitkiewicz, M. Wojnarowska, M. Sołtysik, S.T. Kenny, R.P. Babu, M. Best, K.E. O’Connor, Robust process for high yield conversion of non-degradable polyethylene to a biodegradable plastic using a chemo-biotechnological approach, Waste Manag. 135 (2021) 60–69. https://doi.org/https://doi.org/10.1016/j.wasman.2021.08.030.

## Notes

### Competing Interest Statement

The authors have declared no competing interest.

