## Supplementary material for "Synthetic bacterial consortium for degradation of plastic pyrolysis oil waste"

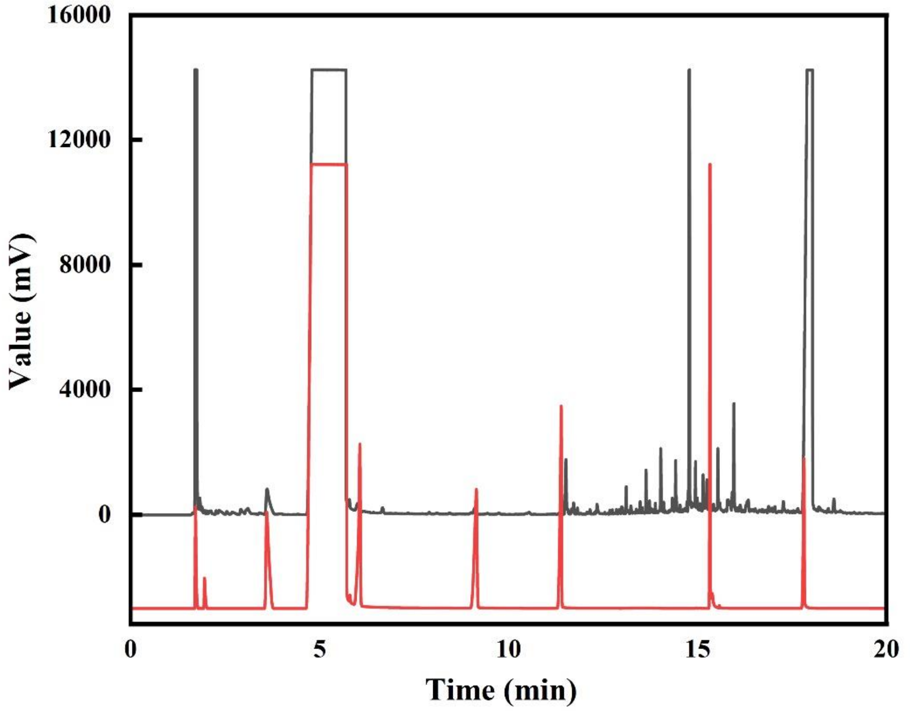


**Fig S1. GC-FID chromatogram of PPOW(black) after filtered and diluted 10 times by n-hexane and standard samples of benzol, toluol, ethylbenzol, cumene, naphthalin and ε-caprolactam(red, from left to right).**

**Benzol**

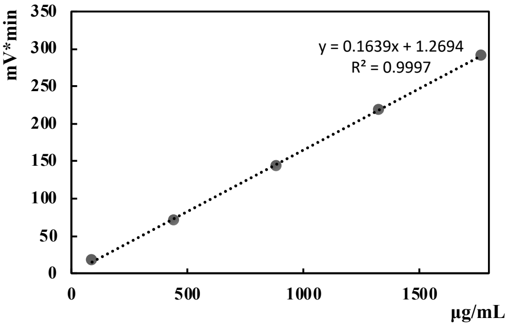


**Ethylbenzol**

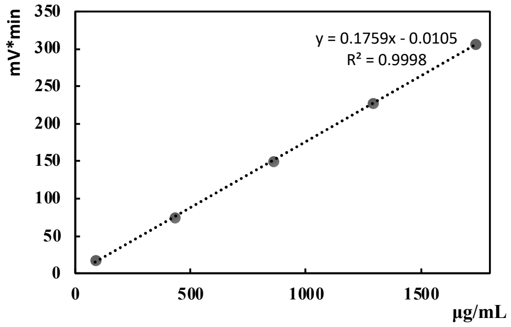


**Toluol**

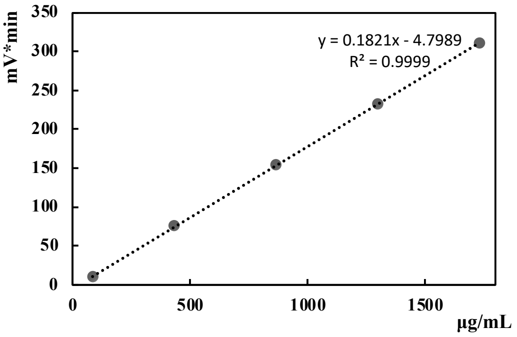


**ε-Caprolactam**

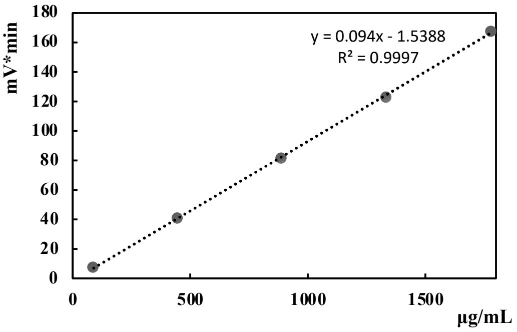

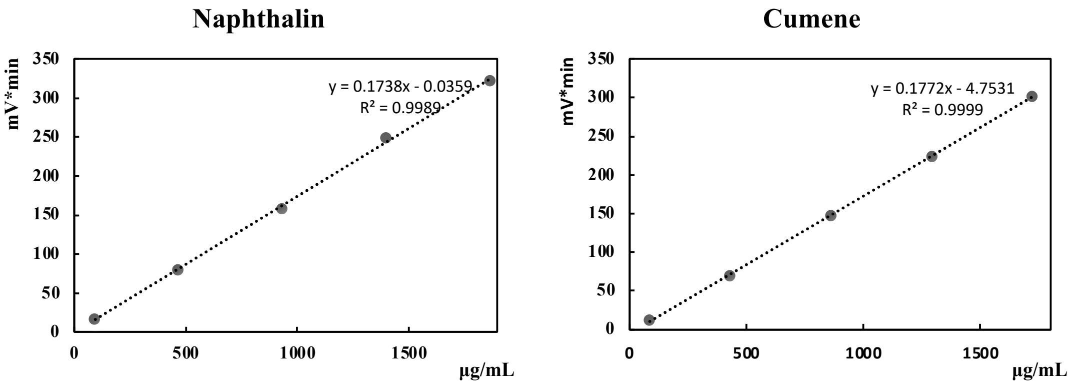


**Fig S2. Cumene,toluol,ethylbenzol, benzol, naphthalin and ε-caprolactam calibration curves by GC-FID.**

**Table S1. Concentrations of several major pollutants other than alkenes in PPOW**

| Name | Cumene | Toluol | Ethylbenzol | Naphthalin | ε-Caprolactam |
|---|---|---|---|---|---|
| Content (μg/mL) | 739.5 | 545.5 | 286.5 | 205.7 | 8032.0 |

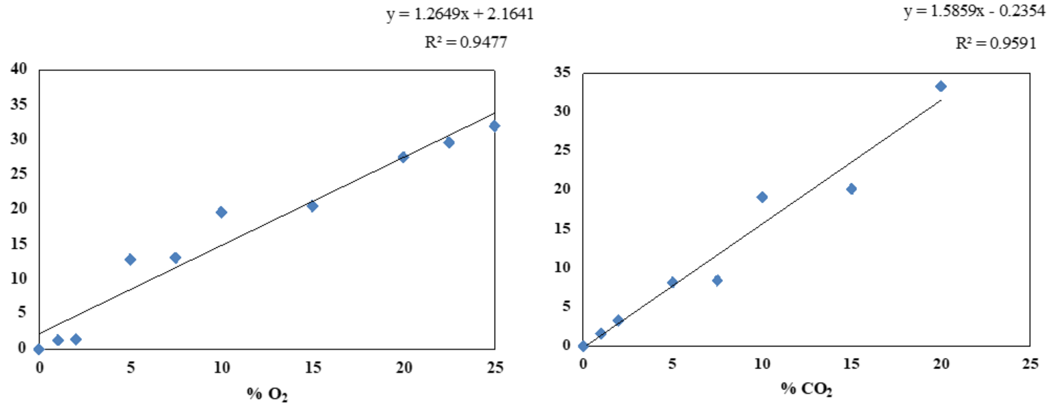


**Fig S3. $CO_2$ and $O_2$ calibration curves by gas chromatograph.**

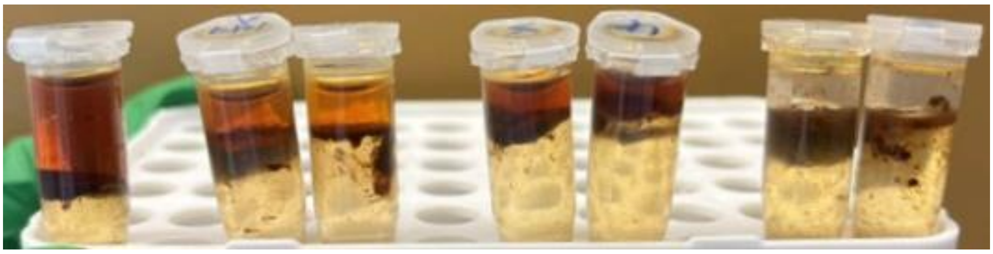

**Fig S4. PPOW after treated by different strains compounding ratios of *Rhodococcus opacus* DSM 43250: *Pseudomonas putida* KT2440: *Pseudomonas spec.* VLB120. From left to right:NC (negative control groups without adding bacteria, 7 days), 1:1:2 (3 days), 1:1:2 (7 days), 2:1:1 (3 days), 2:1:1 (7 days), 1:1:1 (7 days), 1:1:2 (7 days) at 30°C with a second phase–water ratio of 1:5 and 20%( %V) of PPOW content in second phase.**
